# Personalized Metabolite Biomarker Predictions Reveal Heterogeneous Characteristics of Parkinson’s Disease

**DOI:** 10.1101/2025.09.16.676636

**Authors:** Ecehan Abdik, Tunahan Çakır

**Affiliations:** Department of Bioengineering, Gebze Technical University, Kocaeli, Turkey

## Abstract

Understanding the heterogeneous nature of Parkinson’s disease is crucial for improving diagnostic and treatment strategies that benefit distinct patient subgroups. Genome-scale metabolic models, when integrated with omics data, provide powerful frameworks for such investigations. Here, we predicted patient-specific metabolite secretion patterns in the form of oversecretion/undersecretion by the TrAnscriptome-based Metabolite Biomarkers by On–Off Reactions (TAMBOOR) algorithm. We first identified biomarkers for the general PD population using a consensus approach that prioritized changes consistent across the patient cohort. Then, we clustered patients based on the predicted metabolite secretion pattern of each patient to assess heterogeneity and identify potential patient subgroups. Three main clusters were detected, and the most discriminative metabolites underlying this grouping were determined. The power of the discriminative metabolites in grouping PD patients were confirmed with independent validation data to show the reliability and robustness of our approach. Predicted biomarkers for the general population of PD included both well-known disease markers, such as dopamine and eumelanin, and additional metabolites, such as salsolinol, leukotriene A4, heme metabolism products, calcitriol, and retinal, with potential roles in PD mechanism and symptoms. A subset of the predictions also indicated that some well-known characteristics may not be consistently exhibited in all patients. Furthermore, certain metabolites such as melatonin, sphingosine, and biliverdin, though not identified by the general approach, showed distinct secretion patterns across patient clusters. For instance, an undersecretion pattern of melatonin, possibly associated with the sleep disturbance symptom of PD, was detected exclusively in one subgroup. Our study emphasizes the importance of individual-level analysis, which has a high potential to investigate heterogeneity in the disease metabolism. Furthermore, it gives insights into the ways of patient classification that can guide more effective diagnostic and treatment strategies.

## INTRODUCTION

Parkinson’s disease (PD) is a highly prevalent and complex neurodegenerative disease with heterogeneous symptomatology, including clinically important non-motor symptoms such as constipation, depression, and sleep disruption besides motor symptoms (Bloem, Okun, & Klein, 2021; Kalia & Lang, 2015). Although the exact cause of PD is unknown, the complex interaction of genetics and environmental factors contributes to the disease development and progression (Kalia & Lang, 2015). PD also exhibits metabolic complexity, with interconnected metabolic disturbances such as mitochondrial dysfunction, oxidative stress, and impaired protein homeostasis that play critical roles in the disease initiation and progression (Malkus, Tsika, & Ischiropoulos, 2009; Odongo, Bellur, Abdik, & Çak\ir, 2023).

Heterogeneity in the underlying causes and clinical presentation of PD underscores the importance of personalized diagnosis and treatment strategies. Fully individual therapies are not realistic, but certain groups of patients can show better responses to specific therapies (Bloem et al., 2021). Thus, researchers have tried to define subtypes of patients in recent years. Subtype studies commonly depend on clinical symptoms and pathology (Dadu et al., 2022; De Pablo-Fernández, Lees, Holton, & Warner, 2019; Höglinger et al., 2024; Marras & Chaudhuri, 2016). Since molecular-level pathologies emerge before clinical symptoms (Dexter & Jenner, 2013), prediction of molecular biomarkers is crucial for early diagnosis and target-specific therapies. Studies focusing on transcriptomic alterations linked to disease symptoms, progression, and stages have been available in recent years. A study identified differentially expressed genes (DEGs) predictive of AI-defined PD subtypes based on clinical trajectories (Fabrizio, Termine, & Caltagirone, 2025), while another study revealed transcriptomic changes across Lewy Body pathology stages (Cappelletti et al., 2023). Since distinct molecular alterations can lead to same clinical phenotype, identifying molecular alteration-based subtypes in PD is crucial for uncovering subgroups of patients.

Gene expression profiling is the most common strategy for investigating molecular mechanisms of complex diseases that are directly associated with metabolic pathway disruptions, including PD (Kurvits et al., 2021; Zhang, James, Middleton, & Davis, 2005). Integrative usage of genome-scale metabolic models (GEMs) and transcriptome data enables the examination of how changes in gene expression levels impact biological pathways/metabolism (Çak\ir, Abdik, Uzuner, & Lüleci, 2024; Fuhr et al., 2018; Varma et al., 2021; Yizhak, Chaneton, Gottlieb, & Ruppin, 2015). Within the neurodegenerative disease context, this approach was used to identify key metabolic dysregulations as a source of heterogeneity across Alzheimer’s disease (AD) and PD patient subclasses (Lam et al., 2021) and to predict metabolic pathway signatures in PD patient-derived neural precursor cells (Zagare et al., 2023).

This integrative usage is also promising for biomarker prediction by allowing the prediction of oversecretion or undersecretion tendency of metabolites in disease. The prediction of metabolite biomarkers is crucial for advancing clinical diagnostics, as many of these molecules can be non-invasively measured in patient-derived fluids such as blood, urine, saliva, and cerebrospinal fluid. Metabolite biomarker prediction algorithms based on GEMs and transcriptome data were successfully implemented in studies predicting markers for toxicant-induced organ damage in rats (Blais et al., 2017; Pannala et al., 2020; Rawls et al., 2019), experimental mouse models of PD (Abdik & Cakir, 2021), and human brain metabolic changes in PD (Abdik & Çak\ir, 2024). Specifically, TrAnscriptome-based Metabolite Biomarkers by On–Off Reactions (TAMBOOR) algorithm (Abdik & Çak\ir, 2024) accurately predicted aberrant metabolite secretion events in PD.

In this study, the TAMBOOR algorithm was applied to each patient from eight PD-related post-mortem substantia nigra transcriptome datasets obtained from Gene Expression Omnibus (GEO) (Edgar, Domrachev, & Lash, 2002) by integrating the unified human GEM (Human-GEM) (J. L. Robinson et al., 2020) to evaluate the heterogeneous mechanism and pathology of the disease. Predicted metabolite secretion patterns of each of 106 PD patients were examined with a clustering strategy to identify whether subgroups of patients exist in terms of differences in the secretion tendencies for individual metabolites. Then, the most important metabolites for clustering behaviour were identified with feature selection. Another PD dataset, which includes transcriptome data of living brain (LB) samples of the prefrontal cortex (PFC) (de los Santos et al., 2024), was used as the independent validation data to enhance the reliability of our findings. Our biomarker predictions include both hallmarks of disease, like decrease in dopamine and eumelanin secretion, and also other metabolites like leukotriene A4, calcitriol, and retinal, which have important roles in PD-related metabolic pathways. The results also suggest that certain hallmark features of PD are not uniformly presented among patients. Additionally, metabolites such as melatonin, sphingosine, and biliverdin, which were not identified by classical prediction, demonstrate distinct secretion patterns in cluster-based analysis. Our predictions provide molecular-level insight into studies aiming to improve diagnosis and treatment strategies, especially personalized and targeted therapeutic strategies.

## MATERIALS AND METHODS

### Genome-scale Metabolic Model

Human-GEM (version 1.18.0), the generic genome-scale metabolic model of *Homo sapiens*, consisting of 12,995 reactions and 2,899 genes was used in this study. In the simulations, 44 metabolites representing the Ham’s medium and three additional metabolites (NH3, taurine, and ornithine) were used as nutrients to support optimal cellular processes. During simulations, uptake of all metabolites except those 47 metabolites, which are mainly amino acids, vitamins, carbon and oxygen sources, were blocked. The medium metabolites were constrained based on their relative rates reported in the literature as follows: The maximum O2 uptake rate was constrained as 5.5 times the maximum glucose uptake rate (0.32 μmol/g/min) (Çak$ι$r, Alsan, Sayba\cs$ι$l$ι$, Ak$ι$n, & Ülgen, 2007; Sertba\cs, Ülgen, & Çak\ir, 2014), while the maximum uptake rates of amino acids, vitamins and other carbon sources were constrained as 1/10 of the maximum glucose uptake rate (Çak$ι$r et al., 2007). To sustain macromolecule synthesis in all simulations, the minimum biomass production rate was adjusted as 0.0001 μmol/g/min.

### Transcriptome Data

Eight post-mortem brain transcriptome datasets from PD patients were retrieved from Gene Expression Omnibus (https://www.ncbi.nlm.nih.gov/geo/) based on the following criteria: (i) The tissue of origin must be Substantia nigra (SN), (ii) each dataset must include at least 10 patients. The datasets (GSE7621, GSE8397, GSE20141, GSE20292, GSE26927, GSE24378, GSE114517, and GSE49036) (Dijkstra et al., 2015; Durrenberger et al., 2012; Lesnick et al., 2007; Moran et al., 2006; Simchovitz et al., 2020; Zhang et al., 2005; Zheng et al., 2010) were used to predict metabolites with altered secretion tendency between disease and control states. Non-normalized forms of the datasets were obtained from Gene Expression Omnibus. While the microarray datasets were normalized with robust multi-chip average (RMA) and quantile normalization methods, the Trimmed Mean of M-values (TMM) method was applied for the normalization of the RNA-seq dataset as detailed in our previous study (Odongo et al., 2023). PD samples of GSE49036 have Braak stage information, and only those with a Braak stage 3-6 were labelled as PD in this study. 1 sample (GSM663086) from GSE26927, 2 samples (GSM4054769 and GSM4054767) from GSE136666 and 3 samples (GSM606624, GSM606625 and GSM606626) from GSE20292 were identified as outliers based on principal component analysis (PCA) and excluded. Following outlier removal, 106 gene expression profiles of PD patients were retained from the eight datasets, which were used as the main prediction data.

Another dataset, which includes RNA-seq data of living brain (LB) samples from the prefrontal cortex (PFC), was used as validation data. The LB samples were collected from individuals undergoing deep brain stimulation electrode implantation surgery as a part of the Living Brain Project study (available in www.synapse.org with Synapse ID: syn26337520) (de los Santos et al., 2024). The LB data was filtered to include only samples from individuals aged 60+ and samples from the left hemisphere, where the influence of disease started earlier compared to the right hemisphere. The age criterion mainly depends on the typical diagnosis age of PD. Also, including samples from younger individuals would cause incompatibility with post-mortem data. After data filtration, 81 PD patients and 15 controls were retained in the dataset. PCA was applied to the filtered dataset to check outliers, and 2 PD outlier samples were removed. The TMM method of EdgeR package (M. D. Robinson & Oshlack, 2010) was applied to normalize the data. Since the data was collected and sequenced over an extended period, Surrogate Variable Analysis (SVA) was used to identify unknown variables/covariats in the data (Leek, Johnson, Parker, Jaffe, & Storey, 2012). Then, the effect of identified surrogate variable was removed as a covariate by RemoveBatchEffect, a function of the limma package (Ritchie et al., 2015) in R.

### Personalized Prediction of Metabolite Secretion Tendency by the TAMBOOR Algorithm

The TAMBOOR algorithm is based on mapping expression fold changes from transcriptome data onto the reactions of a human genome-scale metabolic model, followed by the prediction of the secretion tendency of each metabolite for the patient of interest and for the control group (Abdik & Çak\ir, 2024). Metabolites with higher/lower secretion tendency in the patient are considered as candidate biomarkers (oversecreted/undersecreted metabolites). From a technical point of view, the TAMBOOR algorithm basically comprises three steps: (i) calculation of reaction activity changes between two conditions by mapping transcriptome data on the GEM, (ii) calculation of maximum secretion rates of external metabolites by Flux Balance Analysis (FBA), and (iii) prediction of number of active reactions required to support the high secretion rates of metabolites, ensuring consistency with mapped transcriptome data.

In the first step, gene expression values of each PD patient were mapped on Human-GEM through gene–protein-reaction (GPR) associations of reactions in the GEM to define patient-specific reaction scores. The MapExpressionToReactions function from the Cobra Toolbox (Heirendt et al., 2019) was used for mapping in default setting. For the control-state reaction scores, the mean expression values of the control individuals in that dataset were similarly mapped on Human-GEM. The reaction weights were calculated as fold changes of transcriptome-based reaction scores (control/patient for disease state, and patient /control for control state simulations). For the reactions without GPR rules, 1 was assigned for internal reactions and 2 was assigned for external reactions.

As the second step, a linear-programming based optimization was applied to predict the maximum secretion rate of each metabolite in the GEM within the given constraints. In the third step, the secretion rate of a metabolite was constrained to be higher than 90% of the predicted maximum secretion rate (Equation 2). Using this threshold, a secondary optimization predicted reaction routes to sustain high-level metabolite secretion. In this optimization, the calculated reaction weights were used as objective function coefficients to minimize the weighted sum of fluxes (Equation 1). This weight-based minimization prioritized reactions/paths having lower objective function coefficients. Thus, predicted flux distribution reflected the upregulation/downregulation trends. Here, irreversible form of the GEM was used to calculate actual summation of fluxes as detailed elsewhere (Abdik & Çak\ir, 2024; Abdik & Cakir, 2021).

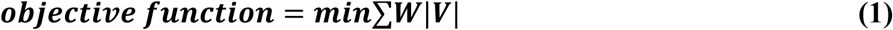

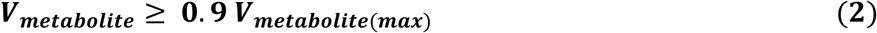

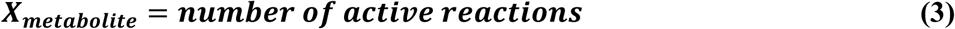

In Equation1-3, W is the weight vector from transcriptome data, V is the flux vector of reactions to be predicted, V_metabolite_ is the secretion rate of the metabolite and X_metabolite_ is the network demand required to support high secretion rate defined in Equation 2. Based on the calculated flux distribution (V), reactions having a flux value higher than 10^-5^ were considered active reactions. For both PD and control cases, the network demand was predicted for the secretion of each metabolite. Then, production score of each metabolite was calculated based on the network demand difference between two conditions as shown in Equation 4.

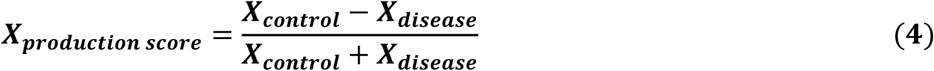

A lower number of active reactions in the disease condition compared to the control indicated a reduced total enzymatic requirement for high-level metabolite secretion. This was interpreted as a greater tendency toward metabolite secretion in the disease state, due to decreased biological resource demand. In this approach, metabolites with the highest production scores were defined as biomarkers with oversecretion potential in the disease case, while those with lowest scores were described as undersecreted metabolites in the disease case. In this study, metabolites found in the upper and lower 15th percentiles of the production score ranking were considered as potentially oversecreted or undersecreted metabolites.

Under the given constraints, the TAMBOOR algorithm generated predictions for 1044 metabolites with defined secretion in Human-GEM, across 106 patients in the main prediction datasets (SN data) and 81 patients in the validation dataset (LB data). Optimization analyses were carried out with MATLAB R2023a and Gurobi 11.0.0.

### Defining Subgroups of Patients by the Clustering Approach

In the second step of TAMBOOR, where maximum secretion rates of metabolites were calculated, 1044 metabolites were detected as produced and released from the Human-Gem under the given constraints. A number of them were dipeptides and tripeptides, with no literature information about their metabolic roles. Therefore, they were excluded from our clustering analysis, which focused on secretion changes and clustering patterns of the remaining 802 metabolites across patients.

TAMBOOR predictions for each patient were coded as 1/-1/0 for metabolites predicted as oversecreted/undersecreted/stable respectively in the patient. In other words, the results were represented as patient-specific categoric vectors, which were combined to create a prediction matrix for each dataset. Patient-specific prediction results of 8 datasets were merged to create a comprehensive prediction matrix including predictions for 802 metabolites and 106 PD postmortem SN samples. The resulting matrix, termed prediction matrix, was used as the main matrix to check if there were any subgroups of patients in terms of predicted biomarkers. The prediction matrix was filtered to exclude features/metabolites with more than 85% of predictions being zero across 106 patients. Then, hierarchical clustering was applied in R using Euclidean distance, and ward.D2 as the linkage method.

After the clustering, features/metabolites important for clustering behaviour were selected by the Random Forest algorithm. The prediction matrix and cluster labels were given to the randomForest function (available under randomForest package) in R to calculate the importance of features. The function calculates importance metrics for each feature. The top 100 most important features based on the importance metric (MeanDecreaseGini) were selected as features important for clustering behaviour. Non-parametric ANOVA (Kruskal-Wallis ANOVA) was applied to check if the selected features were statistically different between clusters, as a validation for the feature selection step (p-value < 0.01).

Based on clusters and selected features of the main prediction matrix, cluster assignment was implemented for the samples of the LB data (the validation data). The K-Nearest Neighbors (KNN) algorithm was utilized to assign a sample to a cluster. The number of neighbours (k) was chosen as 11 based on the square root of the sample size (106 in our case). The distance of an unassigned LB sample to each data point in the prediction matrix was calculated, then the sample was assigned to a cluster by the KNN method based on the majority vote among the k closest neighbours.

To evaluate the biological consistency of the clusters obtained from metabolite-level predictions, transcriptome data of the same samples were used as an independent validation. For consistency between samples from eight different datasets, only the 10,562 genes with matched gene IDs were included in this analysis. Gene expression fold changes were calculated for each sample, and the cluster labels obtained from the metabolite-level predictions were assigned to these expression FC profiles. Expression FC of a gene in a patient was calculated by dividing the gene expression by the mean of the same gene in the control group of a given dataset. The most important 500 genes for the clusters of expression FC profiles were identified by the Random Forest algorithm. Then, enrichment analysis was applied to those genes by the g:Profiler tool (Reimand et al., 2016) to define differential pathways and phenotypes (adjusted p-value<0.05) between clusters.

## RESULTS

Patient-specific metabolite secretion patterns were predicted by the TAMBOOR algorithm to assess heterogeneous metabolic dysregulations in PD. Our assessment proceeded in three stages: (i) identification of metabolite biomarkers for the general PD population based on a consensus approach, (ii) clustering of patients based on individual metabolite secretion patterns and determination of the key discriminative metabolites between clusters, and (iii) validation of both predicted biomarkers and the patient clusters with independent validation data. The overall strategy employed in this study is summarized in Figure 1.

**Figure 1:**
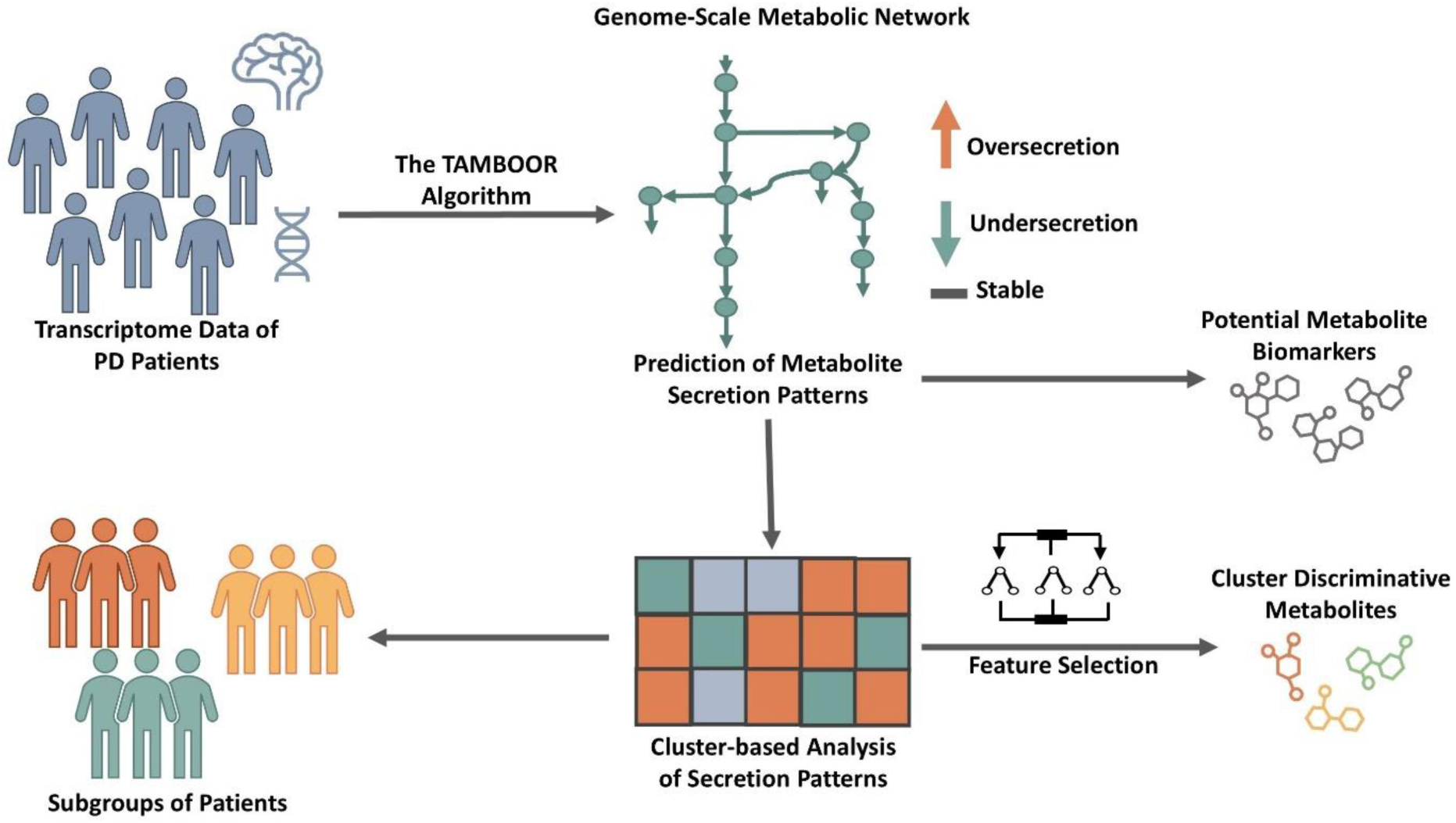
Schematic representation of the general framework of this study. The TAMBOOR algorithm predicts personalized metabolite secretion profiles by combining transcriptome data with a genome-scale metabolic network. Potential biomarkers are predicted based on a majority scoring approach on predicted metabolite secretion patterns. Secretion patterns are subjected to clustering analysis to divide patients into subgroups. Metabolites that are discriminative for clusters are selected by the Random Forest algorithm.

### The TAMBOOR Algorithm Predicts Potential PD Metabolite Biomarkers

Eight transcriptome datasets containing gene expression profiles of 106 PD patients were used as the main data source. For each of the 1044 metabolites with defined secretion in Human-GEM, the TAMBOOR algorithm predicted secretion patterns as oversecreted/undersecreted for each of the 106 patients sourced from independent datasets. After each patient sample was analysed relative to the control group of the corresponding dataset, all personalized predictions were integrated for a comprehensive meta-level evaluation. Considering the heterogeneous characteristics of the disease, metabolites predicted as oversecreted or undersecreted consistently in at least 30% of patients were assumed candidate biomarkers for PD. The number of patients with predictions in the opposite direction was also checked for each metabolite to exclude metabolites with a higher number of opposite-direction predictions. The number of predictions for metabolites suggested as oversecreted/undersecreted are given in Supplementary File 1.

As a result, 151 metabolites, 53 with oversecretion tendency in patients while 98 with undersecretion tendency, were defined as the candidate biomarkers respectively across 106 patients. These metabolites and their associated pathways are listed in Table 1. Dopamine metabolism dysregulation is a well-established hallmark of PD, and in line with this, dopamine-related metabolites were highly represented among the predicted metabolite biomarkers. A detailed pathway association analysis on the metabolites predicted as oversecreted or undersecreted biomarkers in the PD patients revealed most of them to be involved in related or consecutive pathways. Those pathways are phenylalanine, tyrosine, tryptophan and catecholamine metabolisms, lipid metabolism, carbohydrate metabolism, energy metabolisms, glycosaminoglycan metabolism, and amino acid metabolism (Table 1). Besides, metabolites belonging to glutathione, vitamins, and protein metabolisms constituted a significant part of the metabolites predicted as undersecreted in PD.

**Table 1:** Metabolites predicted as oversecreted and undersecreted in PD patients and their involved pathways. Bolded metabolites were validated with the validation data (see the relevant section below).

| Pathway | Undersecreted Metabolites | Oversecreted Metabolites |
| --- | --- | --- |
| Phenylalanine/Tyrosine/Tryptophan Metabolism | phenylacetate, **Phenylacetyl**glycine phacgly, **phenylacetyl**glutamine, **phenylacetyl**glycine, xanthurenate, 2,5-dihydroxybenzoate, 5-hydroxyindoleacetate, homogentisate, 4-hydroxyphenylpyruvate, N-methylsalsolinol, **dopamine**, 5-S-Cysteinyldopamine, N-Acetylvanilalanine, 4-hydroxy-2, quinolinecarboxylic acid (Kynurenic acid), quinonoid dihydrobiopterin, 4-coumarate, 5-S-Cysteinyglycine Dopa, 5-S-cysteinyldopa, eumelanin, 4-S-Glutathionyl-5,6-Dihydroxyindoline | salsolinol, 1,2-dehydrosalsolinol, N-Acetyl-Tyrosine, **phenyllactate**, **noradrenaline**, 3,4-**Dihydroxyphenylethanol**, 4-hydroxyphenyllactate, 3,4-dihydroxyphenylethyleneglycol, dopamine |
| **Lipid Metabolism** | **dihomo-gamma-linolenate, 3-HydroxyisovalericAcid, adrenic acid, (4Z,7Z,10Z,13Z,16Z)-DPA, palmitolate, (R)-mevalonate, DHA, (9E)-tetradecenoic acid, Methyl-Succinate, lathosterol, LPL (Lipoprotein Lipase), Ethylmalonic Acid, (Z,Z,Z)-7,10,13-Hexadecatrienoic Acid, (9Z,12Z,15Z,18Z,21Z)-TPA (6Z,9Z,12Z,15Z,18Z)-TPA, 12,15,18,21-tetracosatetraenoic acid, (6Z,9Z,12Z,15Z,18Z,21Z)-THA, (9Z,12Z,15Z,18Z)-TTA, 1,2-diacylglycerol-LD-, TAG pool, Sebacyl Carnitine** | **leukotriene A4, 5,12,20-TriHETE, GM1b, 6-trans-LTB4, 4-hydroxy-2-nonenal, nonanoic_acid, 2-arachidonoylglycerol, VLDL remnant, HDL, sialyl(2,3)-sialyl(2,6)-galactosylgloboside, glycerol, Suberyl Carnitine, physeteric acid, (7Z)-tetradecenoic acid, 10,13,16-docosatriynoic acid, 9-O-acetylated-GD3, sn-glycerol-3-PC** |
| **Carbohydrate/Energy Metabolism** | **mannose, formate, D-Arabitol, glycerone, glyceraldehyde, glycogenin G4G4, glycogen, galactose, lactose, D-xylulose, ribose, oxalate, glyoxalate, glycolate, D-gluconic acid** | **glucose, pyruvate, glucose-1,6-bisphosphate, fructose, AKG, fructose-1,6-bisphosphate, cis-aconitate, succinate semialdehyde, DHAP, glyceraldehyde** |
| **Iron/Hemoglobin/Heme Metabolism** | **Fe3+, haptoglobin, prothrombin, bilirubin** | **heme, bilirubin-bisglucuronoside, bilirubin-monoglucuronoside, protoporphyrin** |
| **Glycosaminoglycan Metabolism** | **keratan sulfate I, hyaluronate, hyaluronan biosynthesis precursor 1, keratan sulfate I degradation product 1** | **heparan sulfate proteoglycan, UDP-glucuronate, N-acetylneuraminate, N-acetylgalactosamine** |
| **Amino Acid Metabolism** | **saccharopine, homocarnosine, homocysteine, carnosine, Glutaconate, S-Sulfo-L-Cysteine** | **3-Methyl-Glutarate, [protein]-L-lysine, homocarnosine** |
| **Branched-Chain Amino Acid Metabolism** | **2-methylbutyrylglycine, isovalerylglycine, isobutyrylglycine, 3-hydroxyisobutyrate, 2-Methyl-3-Hydroxy-Butyrate, 2-Hydroxy-Isovalerate, 3-methyl-2-oxobutyrate, 2-oxobutyrate, 4-methyl-2-oxopentanoate** | **2-Hydroxy-3-Methyl-Valerate** |
| **Glutathione metabolism** | **GSSG (Glutathione disulfide), GSH, Cys-Gly (Cysteinyglycine)** |  |
| **Vitamin Metabolism** | **vitamin_D3, retinal, (24R)-24,25-dihydroxycalcitol, calcitriol** |  |
| **Proteins/ Protein Metabolism** | **albumin, antitrypsin, [apotransferin], antichymotrypsin, apoB100, apoE, apoC2, apoC1, apoA1** |  |
| **Others** | **H2O2, hydroxide, Ppi, de-Fuc form of PA6 (wo peptide linkage)** | **H+, deoxyribose, N-acetyl-D-mannosamine, PA6, inositol-1-phosphate** |

The most common hallmarks of PD, reduction in dopamine and eumelanin levels, were predicted in at least 30% of patients. On the other hand, some conflicting results were observed regarding the compatibility with the literature and consistency between patients. For example, metabolites such as glyceraldehyde, homocarnosine, and dopamine were predicted to be both oversecreted and undersecreted in an equal number of patients. Moreover, certain metabolites predicted to be oversecreted (e.g., inositol-1-phosphate, HDL, N-acetylneuraminate) were also predicted to be undersecreted in a substantial proportion of patients, while some metabolites predicted as undersecreted (e.g., N-methylsalsolinol, adrenic acid, hydroxide) were likewise predicted to be oversecreted in a considerable number of patients (Supplementary File 1). These findings highlight the heterogeneous characteristics of the disease. Therefore, it is crucial to examine patients individually, as some may not exhibit all typical disease hallmarks. Additionally, investigating personalized therapeutic approaches could benefit specific patient subgroups with distinct responses. It is also crucial to assess predictions without restricting them by the direction of change since secretion patterns of metabolites may diverge across patients with different disease subtypes.

### Clustering of Predicted Metabolite Secretion Behaviours Reveals Subgroups of PD Patients

Investigating the potential of subgrouping arising from the heterogeneity of the disease could be possible with a directionally constraint-free approach. The undersecretion/oversecretion behaviour of a metabolite can exhibit inverse trends between patient groups. Thus, hierarchical clustering was applied to the prediction matrix derived from the SN transcriptome data to investigate potential patient subgroups based on predicted metabolite secretion patterns. This matrix included 595 metabolites (remaining after filtering) across 106 patients, where secretion patterns were encoded as 1 (oversecretion), -1 (undersecretion), and 0 (stable) for each patient. The results were visualized as a heatmap (Supplementary Figure 1). Three main clusters were identified. 106 patients in the cohort were distributed to the three clusters as 34, 37 and 35 patients.

Among the 100 most discriminative metabolites/features identified by the Random Forest algorithm across clusters, 98 showed also statistically significant differences between clusters (Kruskal-Wallis ANOVA, p < 0.01). The defined clusters and the top 100 most important metabolites separating them, out of 595 metabolites, are visualized as a heatmap in Figure 2a. Sixty percent of those important metabolites aligned with the candidate potential biomarkers predicted in the previous section, including dopamine, H_2_O_2_, 4-coumarate, apoA1 and adrenic acid. While certain metabolites like melatonin, prostaglandins, sphingosine, biliverdin, tyramine, and betaine etc., which are associated with PD or neurodegeneration, were not predicted as potential biomarkers in the classical approach, they demonstrated group-specific patterns in the cluster-based approach. The identified metabolites are from diverse categories such as central carbon metabolites, lipids, and amino acid derivatives. The identified discriminative features successfully grouped patients from 8 different datasets into the three clusters without any dataset-specific bias, demonstrating the strength of our approach. Cluster C1 included patients from 8 datasets, C2 included patients from 6 datasets, and there were patients from 7 datasets in Cluster C3.

**Figure 2:**
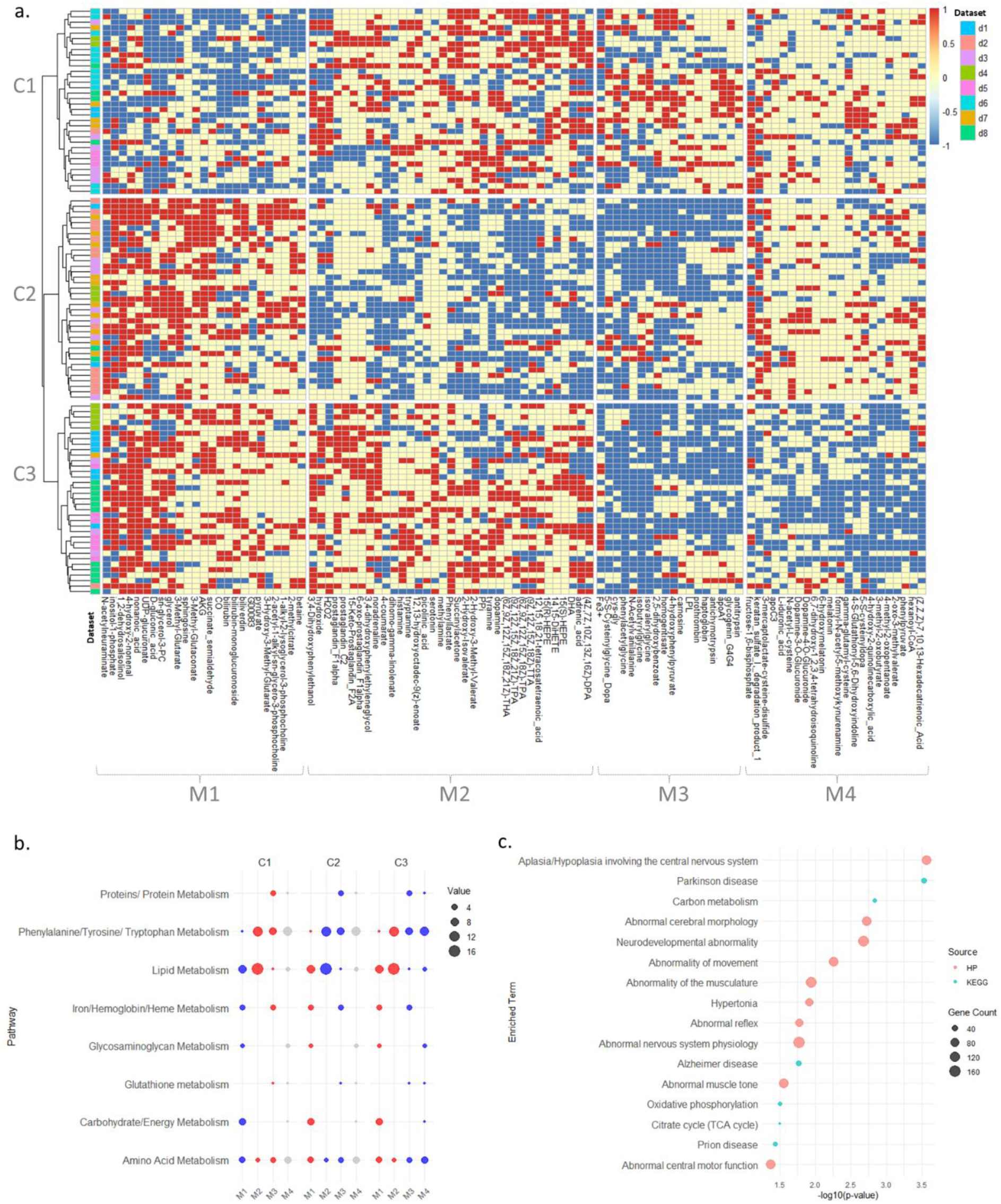
**a)** Heatmap of 100 discriminative features (metabolites) derived from SN transcriptome data based on oversecretion/undersecretion patterns across 106 patients (C1, C2, C3: patient clusters, M1, M2, M3, M4: metabolite clusters). **b)** Pathways of the metabolites included in each metabolite cluster and how they change in patient clusters (blue: undersecretion, red: oversecretion, grey: unchanged/mixed behaviour). **c)** Enrichment analysis result of important genes between clusters (HP: human phenotype).

Metabolites/features were also clustered into 4 main groups to identify metabolites with opposing secretion patterns across clusters (Figure 2a). For example, the metabolites inositol-1-phosphate, 1,2-dehydrosalsolinol and AKG (alpha-ketoglutarate) show oversecretion in most patients from clusters C2 and C3 compared to controls, but undersecretion in cluster C1. On the other hand, metabolites homogentisate, apoA1 and prothrombin show an opposite pattern; undersecretion dominant pattern in C2 and C3 and oversecretion in C1. Additionally, some metabolites do not display a consistent secretion pattern across all clusters. For instance, melatonin, 6-hydroxymelatonin, and L-iduronic acid exhibit an undersecretion pattern in cluster C3, while no clear trend is observed in clusters C1 and C2.

Predicted metabolite features were also systematically assessed in terms of metabolic pathways by identifying their associated pathways and determining pathway dominance through metabolite counts belonging to each pathway. The number of metabolites belonging to each pathway in the metabolite clusters and how they change in each patient cluster is presented in Figure 2b. Examination of important features in terms of metabolic pathways revealed that a considerable portion of metabolites that significantly differ between patient groups are related to dopamine/tyrosine/phenylalanine metabolism, lipid metabolism, and amino acid metabolism (Figure 2b). Interesting patterns were also observed, supporting our hypothesis that the secretion behavior of metabolites within a pathway can be completely opposite across different patient subgroups. Metabolites of carbohydrate/energy metabolism had undersecretion pattern in the patients from cluster C1, while they had oversecretion pattern in the other clusters (Figure 2b). More interestingly, in some cases, metabolites of the same pathway within the same cluster showed opposing secretion patterns. For example, some metabolites of lipid metabolism were oversecreted in C1 while the others were undersecreted; and this pattern was reversed in the patients from C2. Through this parallel evaluation of pathway representations and their subgroup-wise changes, our approach deciphered disease heterogeneity through metabolic mechanisms that distinguish patient subgroups, offering potential clinical insights.

In order to provide functional validation of the identified clusters based on the predicted metabolite secretion patterns, the transcriptome-level changes of the patients in the same clusters were also examined. Compared to our advanced bioinformatic approach that transforms gene expression information into metabolite secretion patterns through metabolic networks, this is only based on the expression FCs of patients across the identified clusters. The most discriminative 500 genes that represent the variation of expression FC values across the clusters were identified. Enrichment analysis was applied to the discriminative genes to observe their relevance to PD. Here, we hypothesize that if the three clusters identified based on predicted metabolite secretion patterns are PD-relevant, the same patients will also show discriminative expression FC patterns in PD-related genes across the clusters. The results revealed that the discriminative genes are enriched in the functional terms directly related to PD (Figure 2c). The enriched KEGG pathways are Parkinson disease, carbon metabolism, Alzheimer disease, oxidative phosphorylation, citrate cycle (TCA cycle) and Prion disease. Enriched human phenotype terms are also highly associated with PD phenotypes such as abnormality of movement, abnormality of the musculature, hypertonia, abnormal reflex, abnormal muscle tone, abnormal central motor function.

### An Independent Dataset Validates Biomarkers Predictions and Clustering Behaviours

Predicted metabolite secretion patterns and their clustering behaviors were also validated by applying the TAMBOOR algorithm on an independent PD transcriptome data, from the Living Brain Project. Almost half of the potential PD biomarkers (26 oversecreted and 43 undersecreted metabolites) predicted by the SN data from the eight datasets were validated with the LB data (Table 1). Identified metabolites spanned nearly all dysregulated pathways listed in Table 1, highlighting the robustness of our approach. The validated metabolites dopamine and salsolinol are involved in dopamine metabolism, and their altered secretion patterns - dopamine undersecretion and salsolinol oversecretion - are well-known metabolic changes in PD. Validation of undersecreted vitamin metabolites vitamin D3, retinal, and (24R)-24,25-dihydroxycalciol highlights another hallmark of PD, vitamin deficiency.

The robustness of the clusters derived from the main prediction data (SN data) using an independent cohort was evaluated in two ways: (i) assessing the consistency between the cluster behaviors in the main prediction data and independent validation data, and (ii) testing the ability of the clusters and discriminative metabolites obtained from the main prediction data to assign group labels to independent samples. In the initial validation phase, the clustering analysis was repeated on the validation cohort using predicted metabolite secretion patterns for 81 PD samples from the LB data to confirm the robustness of patient classification. This independent analysis detected the same number of clusters as derived from the main prediction data. Thus, the three-group patient classification strategy was validated. Furthermore, the robustness of the 100 most discriminative metabolites, identified from the SN cluster analysis and feature selection, was validated by testing their ability to distinguish between clusters in the LB cohort. Each metabolite was subjected to a Kruskal-Wallis ANOVA test to assess its statistical significance among the LB clusters. This analysis demonstrated that 59% (59/100) of the features were significantly differential (p < 0.01) between the LB clusters, underscoring the robustness of the discriminative metabolites. These validated features include many PD-related metabolites such as dopamine, ROS, and melatonin.

In the second validation phase, cluster labels were assigned to each sample of the LB data with the KNN algorithm depending on the clusters and important features of the SN data (100 metabolites in Figure 2). The cluster assignment classified the samples of the validation cohort (n=81) into three groups as follows: 24 samples (∼30%) were assigned to cluster 1, 32 (∼40%) to cluster 2, and 25 (∼40%) to cluster 3. To visually validate the cluster assignment success, prediction matrices of SN and LB data corresponding to 100 discriminative metabolites were merged into a single matrix and subjected to an independent clustering without effects of predefined labels (Figure 3a). The results demonstrate high concordance with the original labels. In particular, 94.8 % of samples originally assigned to cluster 1, 88.4 % to cluster 2, and 78.3 % to cluster 3 re-clustered together. Overall, 87.2% of all samples maintained their original cluster labels, indicating robust patient classification. The resulting dendrogram also demonstrated that samples from different source datasets have an interspersed distribution, not grouped by source. This indicates the absence of a significant batch effect and confirms that the clusters reflect metabolite secretion patterns. Distribution of the original clusters across the newly identified clusters was also represented by the pie charts in Figure 3b.

**Figure 3:**
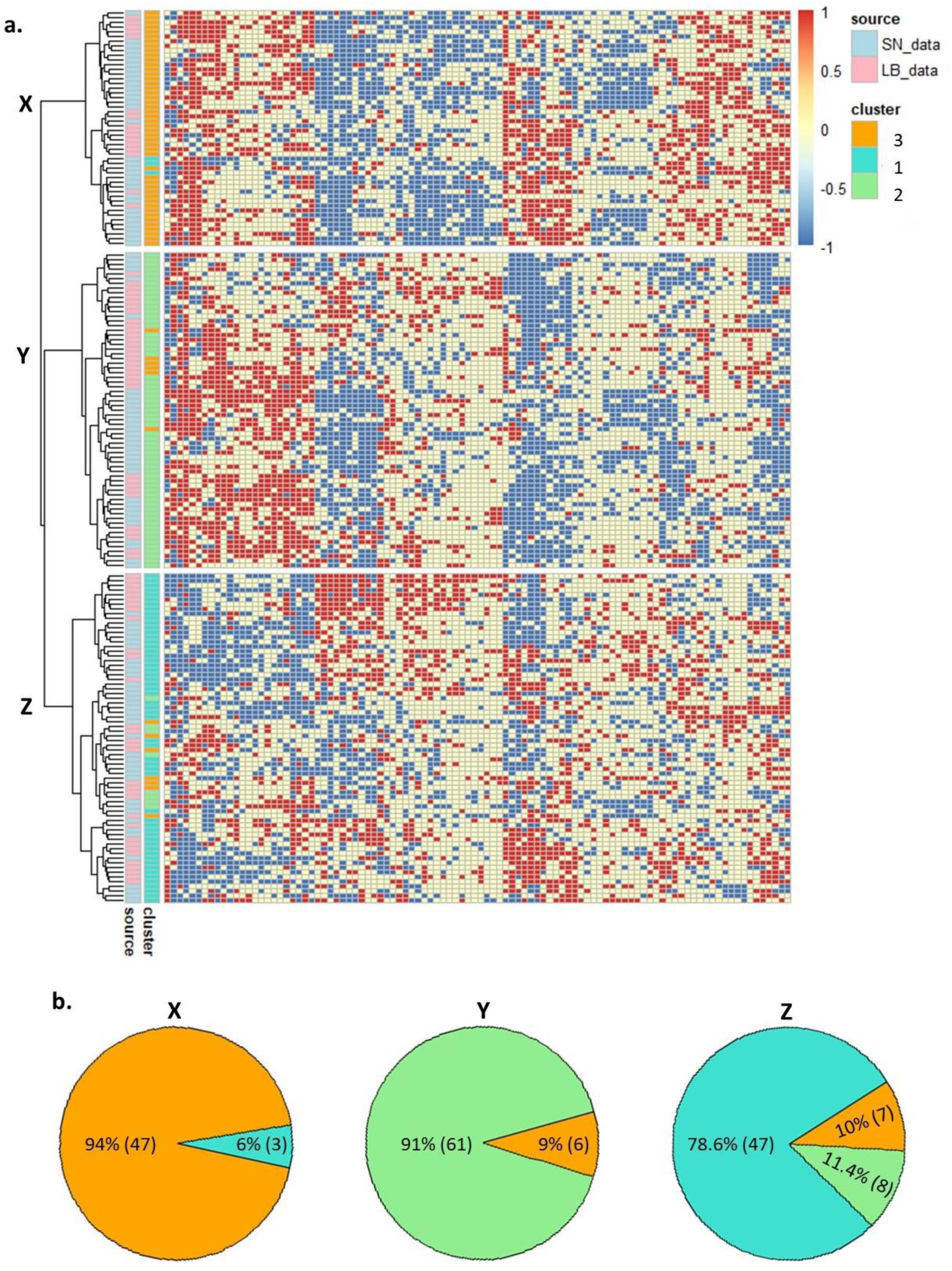
**a)** Unsupervised clustering and a heatmap of the top 100 discriminative metabolites for the combined SN and LB cohorts. **b)** Pie charts display the proportion (percentage) and absolute count of samples from each original cluster (1, 2, 3) within each new cluster (X, Y, Z).

## DISCUSSION

### Predicted Metabolite Biomarkers are Relevant to PD Mechanisms and Pathology

The predicted metabolite biomarkers in this study highlight key metabolic alterations (Table 1) potentially relevant to PD pathology. Phenylalanine, tyrosine, and tryptophan metabolisms have critical roles in maintaining neurological functions, especially through producing neurotransmitters like dopamine depleted in PD (Meiser, Weindl, & Hiller, 2013). Dysfunction of such metabolisms may lead to an imbalance where some metabolites accumulate excessively while others are depleted. Salsolinol, a dopamine-derived neurotoxin, was predicted to be oversecreted for most samples of two datasets (Supplementary File 1 and Table1). Elevated levels of salsolinol were also reported for PD patients with dementia (Antkiewicz-Michaluk, Krygowska-Wajs, Szczudlik, Romańska, & Vetulani, 1997; Voon et al., 2020). Besides dopaminergic neuron loss, its elevated level was suggested as suspicious for visual hallucinations in PD (Moser, Siebecker, Vieregge, Jaskowski, & Kömpf, 1996). Neuromelanin, a dark pigment, is reduced with dopaminergic neuron loss in PD (Poewe et al., 2017). In line with this, our predictions projected a decline in the secretion of dopamine and neuromelanin precursors, such as 5-S-cysteinyldopa and eumelanin. Nevertheless, an increase in dopamine secretion was also predicted in about 30% of patients. This inconsistency likely comes from treatment strategies aiming for dopamine level elevation taken by patients. Another tyrosine metabolism metabolite, 4-coumarate (4-Hydroxycinnamic acid), which was estimated to be undersecreted, has been proposed to be a therapeutic candidate with its neuroprotective effects that can reduce oxidative stress, adjust mitochondrial function, and prevent protein accumulation (Medvedeva et al., 2022; Park et al., 2023).

Within the central nervous system, amino acids have critical roles like being neurotransmitters, neuromodulators, and regulators for energy metabolism. Phenylalanine, tyrosine, and tryptophan are also amino acids directly related with PD (Figura et al., 2018). Amino acids represent a major fraction of differentially abundant metabolites in PD. However, reported level changes are highly inconsistent among studies (Jiménez-Jiménez, Alonso-Navarro, Garc\’\ia-Mart\’\in, & Agúndez, 2020). Differences in dietary/protein consumption of patients and the gut microbiome imbalances commonly observed in PD patients are the most probable cause of this variability (G\katarek et al., 2022). Carnosine, a dipeptide molecule consisting of β-alanine and L-histidine, has crucial roles in muscles and the brain, which are tissues exhibiting high oxidative metabolism. Its reduction causes loss of neuroprotective effects in neurodegenerative diseases (Solana-Manrique, Sanz, Mart\’\inez-Carrión, & Paricio, 2022). The antioxidant power of carnosine has been studied to treat PD (Zhao, Shi, & Zhang, 2017). Predicting carnosine and its derivative homocarnosine as undersecreted metabolites in PD is coherent with this information and our previous study (Abdik & Çak\ir, 2024). Although experimental studies (Mally, Szalai, & Stone, 1997; Molina et al., 1997) and our previous predictions (Abdik & Çak\ir, 2024) supported that lysine level is reduced in PD, lysine secretion was predicted to be increased in about 35% of PD patients.

Impairments in glucose metabolism/carbohydrate metabolism induce energy metabolism dysfunctions, oxidative stress, and implicitly neuroinflammation in PD. Insulin resistance has been associated with PD, especially severe types of PD such as PD with dementia. Hyperglycemia promotes PD-related impairments, including dopamine mechanism dysfunctions, energy metabolism deficiency, and neuroinflammation (Dai et al., 2023). An increase in other sugar molecules, fructose and mannose, in PD was also reported (Trezzi et al., 2017). Nevertheless, a conflicting study that address reduced levels of glucose and mannose in PD is also available (Öhman & Forsgren, 2015). Consistent with our prediction, the end product of glycolysis, pyruvate, was found to be elevated in the plasma of PD patients (Ahmed, Santosh, Kumar, & Christlet, 2009). Reduction in oxidative decarboxylation of pyruvate resulting from deficit mitochondrial respiration may be the main cause of pyruvate accumulation (Mytilineou et al., 1994). An opposing study that reported reduced pyruvate levels in PD is also available (Wuolikainen et al., 2016). Due to pentose phosphate pathways being responsible for the production of lipid metabolism cofactors, carbohydrate metabolism impairments may also be responsible for lipid metabolism impairments in PD (Dai et al., 2023). Formate was predicted to be undersecreted in more than 30% of patients, differing from both our previous prediction study and the literature (Abdik & Çak\ir, 2024; Lock, Zhang, & Checkoway, 2013).

Lipid metabolism dysregulations, including alterations in different classes of lipid-related metabolites like triglycerides, phospholipids, and lipoproteins etc. have contributing roles in neurodegeneration (Xicoy, Wieringa, & Martens, 2019). Leukotrienes are inflammatory lipid mediators, and they are suggested as contributors for neuronal cell death and protein aggregates in neurodegenerative disease (Strempfl et al., 2022). Inhibition of their rate-limiting synthesis enzyme 5-lipoxygenase (5-lox) has been tested as a target-based treatment strategy to reduce dopaminergic neuron loss and α-synuclein aggregation in PD (Kang, Liou, Hour, Liou, & Fu, 2013; Marschallinger et al., 2020). Consistent with literature, levels of leukotriene-related metabolites, leukotriene A4, 5,12,20-TriHETE and 6-trans-LTB4, were suggested as oversecreted metabolites in our predictions. Lathosterol is a key cholesterol precursor whose level was reported as reduced in PD (Cheng et al., 2011), and also predicted to be undersecreted in this study. Disruptions in lipid/cholesterol metabolism affect structural integrity, signaling, and synaptic functions in neurodegenerative diseases. Lathosterol was proposed as a potential modulator against α-synuclein aggregation in an experimental study (Singh et al., 2025). HDL (High Density Lipoprotein), a cholesterol metabolite, secretion was estimated to be elevated in almost half of the patients. However, a detailed analysis of the prediction counts revealed that it was also predicted to be undersecreted in 30% of patients. Conflicting studies reporting elevated and reduced HDL levels in PD are also available in the literature, supporting our sub-typing based approach (M.-X. Dong, Wei, & Hu, 2021; Xicoy et al., 2019).

Blood-related metabolisms like iron, hemoglobin, and heme metabolisms include a group of metabolites predicted to be altered secretion in PD. Heme, bilirubin-bisglucuronoside, bilirubin-monoglucuronoside and protoporphyrin are metabolites involved in heme metabolism, which is highly related to PD pathogenesis. Increased secretion potential of those metabolites was validated with the validation data. Although studies have reported that heme level can be reduced due to elevated expression of Heme Oxygenase-1 (HO-1), an enzyme metabolizing heme, it is also known that the extracellular heme promotes oxidative stress and neuroinflammation (Campomayor, Kim, & Kim, 2025; Sun et al., 2021). Predicting heme and its degradation products as oversecreted biomarkers indicated that overexpression of HO-1 may be consequent of excessive heme production. Other PD pathological features like mitochondrial dysfunction and Lewy body accumulation are also associated with heme and iron accumulation (Freed & Chakrabarti, 2016; Ostrerova-Golts et al., 2000). The blood proteins, apotransferin, albumin, haptoglobin, and prothrombin, were predicted to be undersecreted in PD. Prothrombin (active form: thrombin) has an essential role in blood coagulation. Many evidences linking dysregulated coagulation to PD pathophysiology have been summarised in the literature (X. Wang et al., 2024).

The dysregulation of glycosaminoglycan metabolism in PD contributes to lysosomal impairments and α-synuclein aggregation (Lehri-Boufala et al., 2015; Raghunathan, Hogan, Labadorf, Myers, & Zaia, 2020). Heparan sulfate proteoglycan, a metabolite of glycosaminoglycan metabolism whose secretion was predicted to be increased, is involved in protein aggregates and mediates cellular uptake of protein fibrils (Ma\“\iza et al., 2018; Raghunathan et al., 2020).

Glutathione (GSH) is a critical antioxidant that is crucial for redox homeostasis. Reduced GSH level contributes oxidative stress, mitochondrial dysfunction and neuronal death in PD (Yan et al., 2025). It was suggested as a potential diagnostic biomarker in different studies (H. Dong et al., 2025; Mischley et al., 2016). Its reduction was also involved in the high-confidence metabolite level changes, reported in our previous study (Abdik & Çak\ir, 2024). Other metabolites predicted to be undersecreted, including albumin, apoA1, 5-hydroxyindoleacetate, calcitriol, and carnosine, also showed secretion behavior consistent with the high-confidence metabolite level changes.

Metabolites related to vitamin D which are vitamin D3, (24R)-24,25-dihydroxycalciol and calcitriol (the active form of vitamin D) and retinal (vitamin A) were predicted to be undersecreted. PD patients exhibited vitamin D deficiency, and calcitriol - the biologically active form of vitamin D - exhibited neuroprotective properties in experimental PD models. Retinal is a carotenoid constituent of visual pigments. PD patients have different visual dysfunctions, which are especially common in patients experiencing hallucinations (Weil et al., 2016). Changes in retinal/retinol (vitamin A) related metabolites are directly related to visual abnormalities. The integration of retinoic acid into novel treatments has been studied to improve therapies (Pareek et al., 2024).

### Metabolic Subgrouping Highlights Pathological Heterogeneity in PD

Although the most common features of PD, such as decreased dopamine and eumelanin secretions, were predicted in at least 30% of patients, several other predictions diverged from experimental literature information or lacked consistency across individuals. This situation emphasizes the need for individual-level assessment and supports the utility of categorizing patients into subgroups with similar biological characteristics. The clustering-based analysis of predicted metabolite secretion patterns revealed distinct patient subgroups (Supplementary Figure 1 and Figure 2a), reflecting the heterogeneous nature of the disease and suggesting that hallmark features of PD are not uniformly present across all individuals. Some of the identified discriminative metabolites for grouping patients were directly related to known metabolic alterations in PD. The treatment strategies for Parkinson’s disease commonly aim to increase dopamine level. L-DOPA administration, MAO-B inhibitors, and COMT inhibitors are the most common therapies that are focused on increasing dopamine production from L-DOPA while inhibiting the breakdown of dopamine (Finberg, 2019). Thus, we concluded that the treatments received by patients and the molecular changes induced by these therapies could be the main reasons for dopamine-related metabolite behaviour changes between clusters. For instance, our predictions demonstrate a correlation between metabolite secretion alterations of dopamine and histamine. This relation is compatible with an experimental study (Yanovsky et al., 2011) focusing on the activation of histaminergic neurons and the potential of dopamine and histamine co-release in the case of L-DOPA administration. In Figure 2a, reactive oxygen species (ROS) like hydroxide and H_2_O_2_ show an oversecretion trend in clusters characterized by elevated dopamine secretion. Induction of oxidative stress is also a side effect of L-DOPA-based treatments (Dorszewska, Prendecki, Lianeri, & Kozubski, 2014; Stansley & Yamamoto, 2013). Tyramine is an amino acid metabolized by monoamine oxidase (MAO). Thus, treatments using MAO-B inhibitors can cause an elevation of tyramine and blood pressure (demarcaida et al., 2006). This can be the reason why dopamine and tyramine showed parallel trends between clusters.

Several PD-related metabolites, though not predicted as general biomarkers by standard prediction assessment, exhibited group-specific secretion patterns in cluster-based analysis. Sleep disturbance is one of the non-motor symptoms seen in PD patients. Melatonin deficiency has been attributed responsibility for this symptom in PD (Ma, Yan, Sun, Jiang, & Zhang, 2022). Most of the patients in cluster 3 (Figure 2a) show undersecretion behaviour in melatonin production. Prostaglandins are mediators in the central nervous system, and changes in their metabolism can contribute to neuroinflammation and neurodegeneration in neurodegenerative diseases, including PD (Lima, Bastos, Limborço-Filho, Fiebich, & De Oliveira, 2012). Inhibition of prostaglandin-produced enzymes has been studied as a treatment strategy (Yagami, Koma, & Yamamoto, 2016). Sphingosine-1-phosphate (S1P), the active form of sphingosine, has crucial neuroprotective roles in PD (W. Wang, Zhao, & Zhu, 2023). However, oxidative stress in PD inhibits the enzyme sphingosine kinases (SphKs) that convert sphingosine to S1P (Motyl, Strosznajder, Wencel, & Strosznajder, 2021). The oversecretion sphingosine pattern observed in the patients in cluster 2 may have resulted from this disturbed sphingosine homeostasis. Carbon monoxide (CO) and biliverdin, which is then converted to bilirubin, are heme degradation products (Hatano, Saiki, Okuzumi, Mohney, & Hattori, 2016). CO, biliverdin, and bilirubin patterns are consistent as oversecreted in clusters 2 and 3 while undersecreted in cluster 1. Betaine has been studied for its protective functions against oxidative stress and neurodegeneration to improve PD treatments and decrease the side effects of L-DOPA administration (Bhatt, Di Iacovo, Romanazzi, Roseti, & Bossi, 2023).

Adrenic acid, a fatty acid, was reported as a promising biomarker and therapeutic target before. Since free adrenic acids promote α-synuclein aggregation, treatment strategies that decrease the level of adrenic acid have been tested to minimize protein accumulation in PD (Z. Wang et al., 2024). Contrary to this literature information, adrenic acid was predicted to be undersecreted in almost half of the patients. Despite this, the cluster-based approach revealed that patients whose adrenic acid secretion tendencies were predicted to be increased clustered together in the patient cluster 3. Picolinic acid is produced from the kynurenine pathway and is known for its neuroprotective role in the central nervous system. Kynurenine pathway alterations are observed in PD pathology (Grant, Coggan, & Smythe, 2009). Thus, picolinic acid is also suspected to be altered in PD. Although its secretion was not predicted to be changed in the majority of prediction, it was detected as metabolite having significantly changed secretion tendency between the clusters.

### Validation Data Confirms Predicted Biomarkers and Patient Grouping Strategy

Assessment of transcriptomic differences across the defined clusters showed that most of the varying genes are linked to PD-related pathways and phenotypes (Figure 2c). These results supported that even the well-known features of the disease are not consistently exhibited by all patients, as mentioned in metabolite-based analysis. In addition to transcriptome-based confirmation, metabolite biomarker prediction with an independent validation dataset verified a substantial portion of predicted biomarkers, including dopamine and numerous metabolites in PD-associated pathways. Almost all of the discriminative metabolites (dopamine, tyramine, ROS, melatonin, biliverdin, bilirubin, CO, betaine, adrenic acid, and picolinic acid) mentioned as having differential secretion patterns that are consistent with the literature above were also confirmed as significantly differential between clusters of the validation data (LB data). The success of cluster assignment for validation cohort samples by metabolites selected from the clustering behaviors of the main prediction data was verified by independent clustering of the merged data including samples of the main dataset and the validation dataset. This analysis demonstrated a homogeneous distribution of samples from the two datasets (Figure 3a) and confirmed that most samples maintained their initial cluster assignments (Figure 3b), thereby validating the robustness of the selected metabolites for grouping PD patients.

## CONCLUSION

Personalized metabolite secretion behavior predictions confirmed the complex and heterogeneous metabolic nature of the disease. Cluster-based analysis of predictions revealed that patient subgroups display contrasting metabolic changes despite common underlying pathways. The significant alterations predicted in PD-related metabolites and genes between subgroups highlighted phenotypic variability and the inadequacy of generalized treatment strategies. Future experimental studies exploring the relationship between these metabolic predictions, disease symptoms, disease severity, and drug responses of patients could make it possible to group patients with group-specific markers. In this way, the highlighted metabolites can gain a high potential for improving personalized treatment strategies and supplement/dietary suggestions with further studies for validation.

