## Supplementary Figure 1 for "Personalized Metabolite Biomarker Predictions Reveal Heterogeneous Characteristics of Parkinson’s Disease"

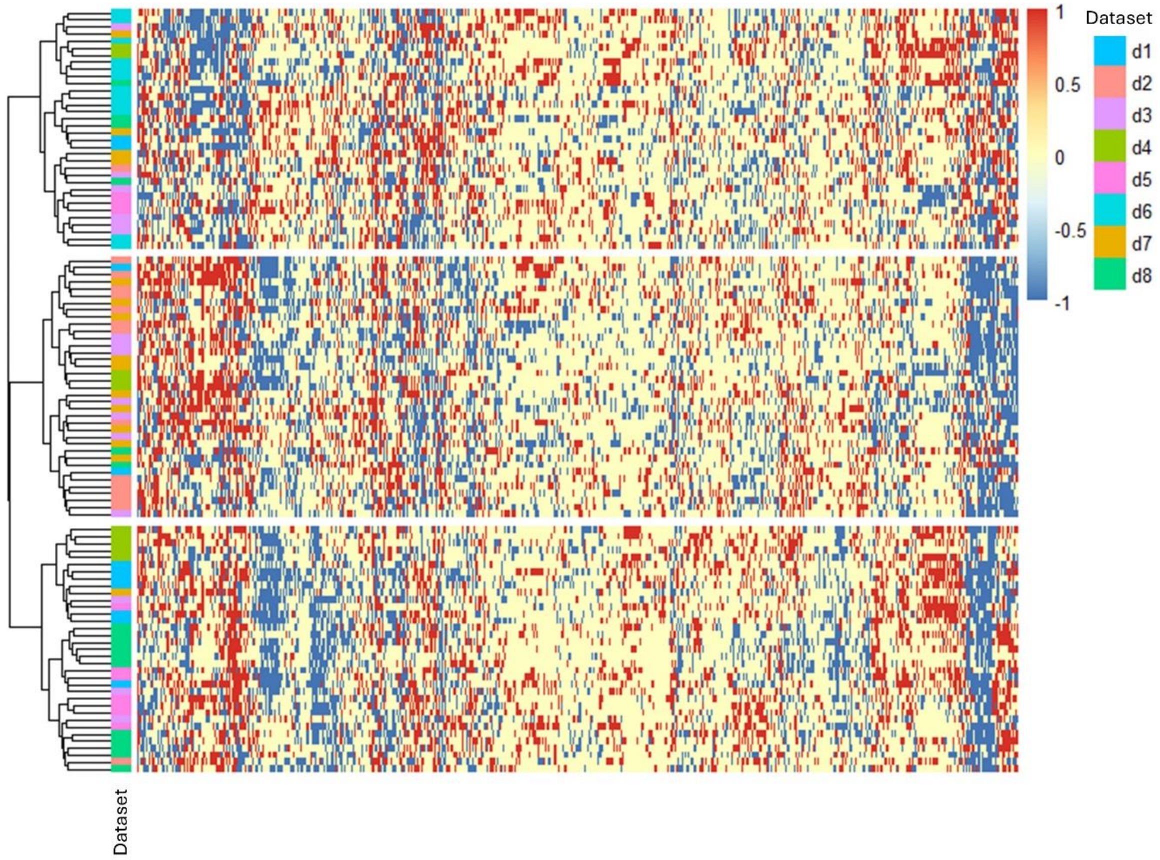


**Supplementary Figure 1:** Heatmap of metabolite secretion patterns across patient samples. The matrix comprises 595 metabolites (columns) and 106 patients (rows). Secretion behaviors are encoded as oversecretion (1), undersecretion (-1), and stable/no prediction (0).
