## Supplementary File 1 for "Personalized Metabolite Biomarker Predictions Reveal Heterogeneous Characteristics of Parkinson’s Disease"

**Supplementary Table 1:** The number of predictions for metabolites suggested as oversecreted/undersecreted.

| **Oversecreted Metabolites** | **Number of Predictions for Oversecretion** | **Number of Predictions for Undersecretion** |
|---|---|---|
| leukotriene A4 | 34 | 7 |
| 4-hydroxy-2-nonenal | 57 | 14 |
| nonanoic acid | 53 | 14 |
| succinate semialdehyde | 36 | 14 |
| cis-aconitate | 33 | 14 |
| H+ | 32 | 14 |
| 6-trans-LTB4 | 32 | 14 |
| VLDL remnant | 40 | 18 |
| AKG | 42 | 19 |
| 3,4-dihydroxyphenylethyleneglycol | 37 | 17 |
| 2-arachidonoylglycerol | 36 | 17 |
| heme | 39 | 19 |
| N-acetyl-D-mannosamine | 41 | 20 |
| 4-hydroxyphenyllactate | 34 | 17 |
| salsolinol | 59 | 31 |
| PA6 | 38 | 20 |
| 3-Methyl-Glutarate | 38 | 20 |
| bilirubin-bisglucuronoside | 39 | 21 |
| [protein]-L-lysine | 38 | 21 |
| glucose-1,6-bisphosphate | 36 | 20 |
| sialyl(2,3)-sialyl(2,6)-galactosylgloboside | 32 | 18 |
| 5,12,20-TriHETE | 35 | 20 |
| N-Acetyl-Tyrosine | 43 | 25 |
| 1,2-dehydrosalsolinol | 50 | 30 |
| glycerol | 44 | 27 |
| N-acetylgalactosamine | 32 | 20 |
| glucose | 41 | 26 |
| inositol-1-phosphate | 52 | 33 |
| 9-O-acetylated-GD3 | 34 | 22 |
| HDL | 49 | 32 |
| GM1b | 32 | 21 |
| pyruvate | 35 | 23 |
| physeteric acid | 38 | 25 |
| heparan sulfate proteoglycan | 44 | 30 |
| deoxyribose | 37 | 27 |
| Suberyl Carnitine | 34 | 25 |
| Phenyllactate | 34 | 25 |
| bilirubin-monoglucuronoside | 40 | 30 |

| | | |
|---|---|---|
| (7Z)-tetradecenoic acid | 37 | 28 |
| protoporphyrin | 33 | 25 |
| noradrenaline | 33 | 28 |
| DHAP | 40 | 34 |
| fructose-1,6-bisphosphate | 40 | 34 |
| 10,13,16-docosatriynoic acid | 36 | 31 |
| fructose | 32 | 28 |
| 3,4-Dihydroxyphenylethanol | 36 | 32 |
| sn-glycerol-3-PC | 39 | 35 |
| UDP-glucuronate | 41 | 38 |
| 2-Hydroxy-3-Methyl-Valerate | 36 | 34 |
| N-acetylneuraminate | 44 | 42 |
| glyceraldehyde | 36 | 36 |
| homocarnosine | 34 | 34 |
| dopamine | 32 | 32 |
| **Undersecreted Metabolites** | | |
| phenylacetylglycine | 15 | 73 |
| 2-methylbutyrylglycine | 10 | 70 |
| isobutyrylglycine | 13 | 68 |
| phenylacetate | 6 | 66 |
| Phenylacetylglycine phacgly | 6 | 64 |
| isovalerylglycine | 12 | 63 |
| phenylacetylglutamine | 11 | 62 |
| 5-S-Cysteinyldopamine | 6 | 61 |
| Fe3+ | 19 | 61 |
| 2,5-dihydroxybenzoate | 18 | 59 |
| cys-gly | 11 | 55 |
| saccharopine | 24 | 55 |
| 4-hydroxyphenylpyruvate | 20 | 54 |
| 3-Hydroxyisovaleric Acid | 20 | 52 |
| N-Acetylvanilalanine | 8 | 50 |
| xanthurenate | 11 | 50 |
| adrenic acid | 32 | 50 |
| N-methylsalsolinol | 33 | 50 |
| 4-coumarate | 14 | 49 |
| homogentisate | 16 | 48 |
| 3-hydroxyisobutyrate | 21 | 48 |
| 4-hydroxy-2-quinolinecarboxylic acid | 8 | 47 |
| 5-S-Cysteinylglycine Dopa | 17 | 46 |
| prothrombin | 10 | 45 |
| H2O2 | 26 | 45 |
| ribose | 29 | 45 |
| D-xylulose | 26 | 44 |

| | | |
|---|---|---|
| 12,15,18,21-tetracosatetraenoic acid | 27 | 44 |
| hyaluronate | 29 | 44 |
| [apotransferin] | 8 | 43 |
| 2-oxobutyrate | 9 | 43 |
| (9Z,12Z,15Z,18Z,21Z)-TPA | 24 | 42 |
| (6Z,9Z,12Z,15Z,18Z)-TPA | 25 | 42 |
| 2-Methyl-3-Hydroxy-Butyrate | 27 | 42 |
| hyaluronan biosynthesis, precursor 1 | 29 | 42 |
| DHA | 32 | 42 |
| dihomo-gamma-linolenate | 11 | 41 |
| antichymotrypsin | 11 | 41 |
| homocysteine | 11 | 41 |
| LPL | 12 | 41 |
| mannose | 19 | 41 |
| (4Z,7Z,10Z,13Z,16Z)-DPA | 27 | 41 |
| haptoglobin | 5 | 40 |
| 3-methyl-2-oxobutyrate | 8 | 40 |
| apoC1 | 10 | 40 |
| apoA1 | 13 | 40 |
| (R)-mevalonate | 23 | 40 |
| palmitolate | 27 | 40 |
| galactose | 30 | 40 |
| de-Fuc form of PA6 (wo peptide linkage) | 31 | 40 |
| D-gluconic acid | 36 | 40 |
| albumin | 7 | 39 |
| (9E)-tetradecenoic acid | 32 | 39 |
| hydroxide | 34 | 39 |
| antitrypsin | 7 | 38 |
| apoC2 | 9 | 38 |
| glycogenin G4G4 | 14 | 38 |
| GSH | 16 | 38 |
| Glutaconate | 20 | 38 |
| lactose | 29 | 38 |
| (9Z,12Z,15Z,18Z)-TTA | 30 | 38 |
| keratan sulfate I, degradation product 1 | 31 | 38 |
| Sebacoyl Carnitine | 32 | 38 |
| apoE | 5 | 37 |
| apoB100 | 8 | 37 |
| (6Z,9Z,12Z,15Z,18Z,21Z)-THA | 23 | 37 |
| (Z,Z,Z)-7,10,13-Hexadecatrienoic Acid | 18 | 36 |
| vitamin D3 | 24 | 36 |
| Methyl-Succinate | 26 | 36 |
| glyceraldehyde | 36 | 36 |

| | | |
|---|---|---|
| S-Sulfo-L-Cysteine | 11 | 35 |
| carnosine | 13 | 35 |
| glyoxalate | 18 | 35 |
| bilirubin | 21 | 35 |
| 1,2-diacylglycerol-LD-TAG pool | 26 | 35 |
| 4-methyl-2-oxopentanoate | 14 | 34 |
| oxalate | 19 | 34 |
| formate | 19 | 34 |
| eumelanin | 20 | 34 |
| (24R)-24,25-dihydroxycalciol | 24 | 34 |
| keratan sulfate I | 28 | 34 |
| D-Arabitol | 28 | 34 |
| glycerone | 31 | 34 |
| homocarnosine | 34 | 34 |
| 5-S-cysteinyldopa | 18 | 33 |
| calcitriol | 19 | 33 |
| glycolate | 20 | 33 |
| 2-Hydroxy-Isovalerate | 23 | 33 |
| glycogen | 25 | 33 |
| quinonoid dihydrobiopterin | 0 | 32 |
| 5-hydroxyindoleacetate | 10 | 32 |
| Ethylmalonic Acid | 11 | 32 |
| PPi | 13 | 32 |
| 4-S-Glutathionyl-5,6-Dihydroxyindoline | 14 | 32 |
| GSSG | 16 | 32 |
| retinal | 23 | 32 |
| lathosterol | 28 | 32 |
| dopamine | 32 | 32 |
